# Pentameric assembly of the Kv2.1 tetramerization domain

**DOI:** 10.1101/2021.11.09.467962

**Authors:** Zhen Xu, Saif Khan, Nicholas J. Schnicker, Sheila A. Baker

**Affiliations:** Protein and Crystallography Facility, University of Iowa, Iowa City, United States; Department of Biochemistry and Molecular Biology, University of Iowa, Iowa City, United States

**Author notes:** Correspondence: Nicholas Schnicker, Protein and Crystallography Facility, University of Iowa, 4-611 BSB, 51 Newton Road, Iowa City, IA 52242, USA., Sheila A. Baker, Department of Biochemistry and Molecular Biology, University of Iowa, 4-712 BSB, 51 Newton Road, Iowa City, IA 52242, USA.

**Keywords:** Voltage-gated potassium channel Kv2.1, tetramerization domain, crystal structure, SAXS pentamer, tetramer

## Abstract

The Kv family of voltage-gated potassium channels regulate neuronal excitability. The biophysical characteristic of Kv channels can be matched to the needs of different neurons by forming homotetrameric or heterotetrameric channels within one of four subfamilies. The cytoplasmic tetramerization (T1) domain plays a major role in dictating the compatibility of different Kv subunits. The only Kv subfamily missing a representative structure of the T1 domain is the Kv2 family. We used X-ray crystallography to solve the structure of the human Kv2.1 T1 domain. The structure is similar to other T1 domains but surprisingly formed a pentamer instead of a tetramer. In solution the Kv2.1 T1 domain also formed a pentamer as determined with in-line SEC-MALS-SAXS and negative stain EM. The Kv2.1 T1-T1 interface involves electrostatic interactions including a salt bridge formed by the negative charges in a previously described CDD motif, and inter-subunit coordination of zinc. We show that zinc binding is important for stability. In conclusion, the Kv2.1 T1 domain behaves differently from the other Kv T1 domains which may reflect the versatility of Kv2.1, the only Kv subfamily that can assemble with the regulatory KvS subunits and scaffold ER-plasma membrane contacts.

## Introduction

Voltage-gated potassium channels (Kv) are essential for regulating membrane potential, propagating action potentials, and controlling potassium homeostasis in a diverse array of neuronal and non-neuronal tissues^1,2^. There are 27 Kv genes in the human genome classified into five subfamilies with distinct expression patterns and biophysical properties: Kv1 (shaker), Kv2 (shab), Kv3 (Shaw), Kv4 (shal), and the regulatory KvS (silent). Functional diversity is achieved by mixing and matching Kv monomers to form heterotetrametric channels in addition to the homotetrameric channels^3^. Only Kv subunits within the same subfamily can interact. The exception to this is the KvS proteins which do not form homomeric channels but instead must assemble with Kv2 subunits^4,5^. The major constraint on inter-subfamily assembly is the cytoplasmic tetramerization domain (T1)^6-9^. How the T1 domain confers selective assembly is poorly understood.

The T1 domain is one of four structural classes of the BTB (or POZ) superfamily of protein-protein interaction domains^10^. The core BTB fold is ∼95 amino acids and comprised of five α-helices organized into two sets of hairpins that is capped at one end by three β-strands. Deletions, insertions, or extensions to the core BTB fold characterizes different structural classes with the T1 class from Kv proteins being the most like the core BTB fold. BTB folds are most often found as homodimers or in the case of the T1 domains, tetramers. One interesting exception is the BTB fold from KCTD proteins which may be monomers, dimers, tetramers or pentamers^11^.

While there are no representative T1 structures from either the Kv2 or KvS subfamilies, structural analysis of T1 domains from Kv1, Kv3, and Kv4 proteins have provided key insights into how selective assembly of tetramers occurs. Some subfamily-specific individual differences in charged or hydrophobic amino acids facing the tetramerization interface have been identified as important for assembly^7,10,12-14^. Another critical factor in selective T1 domain assembly is inter-subunit coordination of zinc. A HX_5_CX_20_CC motif is conserved in Kv2, Kv3, and Kv4 proteins but is absent in Kv1 proteins^12^. In Kv4.2 and Kv3.1 T1 domains Zn^2+^-binding has been shown to be essential for monomers to assemble into tetramers^14-16^. The role of Zn^2+^-binding has not been tested for Kv2 or KvS T1 domains, but it is logical to assume it plays a similar role in assembly. Thus zinc-dependent assembly distinguishes the Kv1 T1 domain from the Kv2, Kv3, and Kv4 T1 domains. A feature that distinguishes Kv2 and KvS T1 domains from Kv3 and Kv4 T1 domains is the presence of a CDD motif^17^. Modeling of the Kv2.1 T1 domain indicated that the critical aspartates in the CDD motif are not at the T1 homotetrameric interface but closer to the surface of the domain. This raises the possibility that an interaction between the surface of Kv T1 domains and other regions of the channel could add to the subfamily-selective assembly dictated by T1 domain tetramerization. Supporting this idea is the observation that the N- and C-termini of Kv2.1 can interact^17-19^. Solving the structures of Kv2 and KvS T1 domains will be necessary steps toward the goal of completing a comprehensive comparative analysis of how T1 domains confer selective assembly.

In this study we solved the crystal structure of the human Kv2.1 T1 domain and investigated the role of Zn^2+^ in the stability of the protein. To our surprise the Kv2.1 T1 domain was pentameric both in crystal and solution. Zn^2+^, along with a salt bridge formed from the aspartates in the CDD motif, provides stability. The different assembly state of the isolated Kv2.1 T1 domain, compared to the isolated T1 domains from Kv1, Kv3, and Kv4 proteins, indicates that *in vivo*, multiple interactions between Kv2.1 subunits are required to ensure proper assembly into tetramers.

## Results

### The crystal structure of the Kv2.1 tetramerization (T1) domain

The T1 domain of voltage-gated potassium channel Kv2.1 (human Kv2.1 T1; residues 29–147) was expressed in *E. coli* BL21 (DE3) and purified by Ni-NTA, followed by TEV cleavage to remove the N-terminal His-tag, and Superdex-200 chromatography. Crystals of Kv2.1 T1 were in space group P4_1_2_1_2. Initial phase estimates were obtained by molecular replacement using a Kv3.1 T1 domain monomer (PDB: 3KVT) as a search model and the resulting Kv2.1 T1 structure was refined at a resolution of 2.5 Å (PDB: 7RE5). Electron densities for residues 29-133 were identified and the missing C-terminal residues (134-147) may have been disordered. Surprisingly, in the Kv2.1 crystals five protein molecules were present in the asymmetric unit, forming a pentameric ring, as shown from the C-terminal (top) and side view (Figure 1A, B). Diffraction data and refinement statistics are presented in Table 1. A metal ion was identified in each of the individual T1 domains, coordinated by a conserved zinc-binding, HX_5_CX_20_CC motif. The Kv2.1 T1 pentameric rings were stacked as dimers in the crystal and closer examination of that interface between rings revealed a second metal ion coordinated by residues introduced by the N-terminal NdeI cloning site. Representative electron densities and verification of the metal as Zn^2+^ by X-ray fluorescence spectroscopy is shown in Figure S1. We also obtained a lower resolution (2.7Å) structure from an alternative space group C222_1_, which was also a pentamer but with partial zinc occupancy (for details see PDB: 7SPD). The core BTB fold of the Kv2.1 T1 monomers is similar to the monomers in the pentameric T1-like domain from KCTD5 (Figure 1D) and tetrameric Kv4.2 T1 domain (Figure 1C). The five Kv2.1 T1 subunits are rotated from the central axis of the ring by 70.5°, 71.3°, 71.5°, 73.2°, and 73.5°, which is more open than the arrangement in KCTD5 (2x 69.5°, 72.8°, 73.4°, and 74.7°). Overall, the pentameric ring of Kv2.1 T1 subunits is less compact than that for KCTD5.

**Figure 1.**
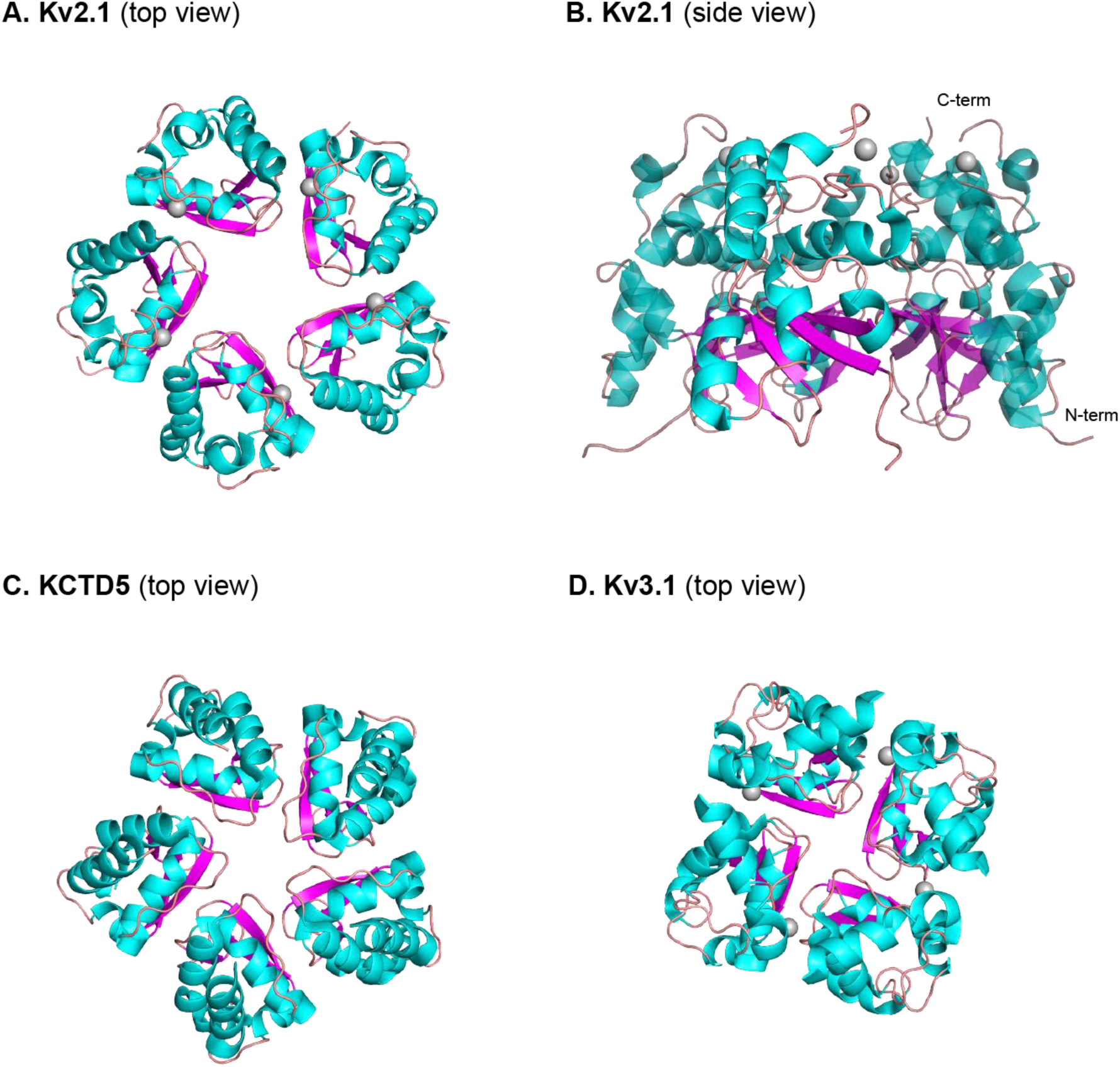
Overview of Kv2.1 T1 crystal structure. **(A)** Cartoon representation of top view (C-terminus) and **(B)** side view of X-ray structure Kv2.1 T1 in pentameric form (7RE5). **(C)** Top view of previously reported X-ray structures for the pentamer of KCTD5 (3DRZ) and **(D)** the tetramer of Kv4.2 (1NN7). Colors for secondary structure are α-helix (cyan) and β-sheet (magenta), the grey sphere is zinc.

**Table 1.**
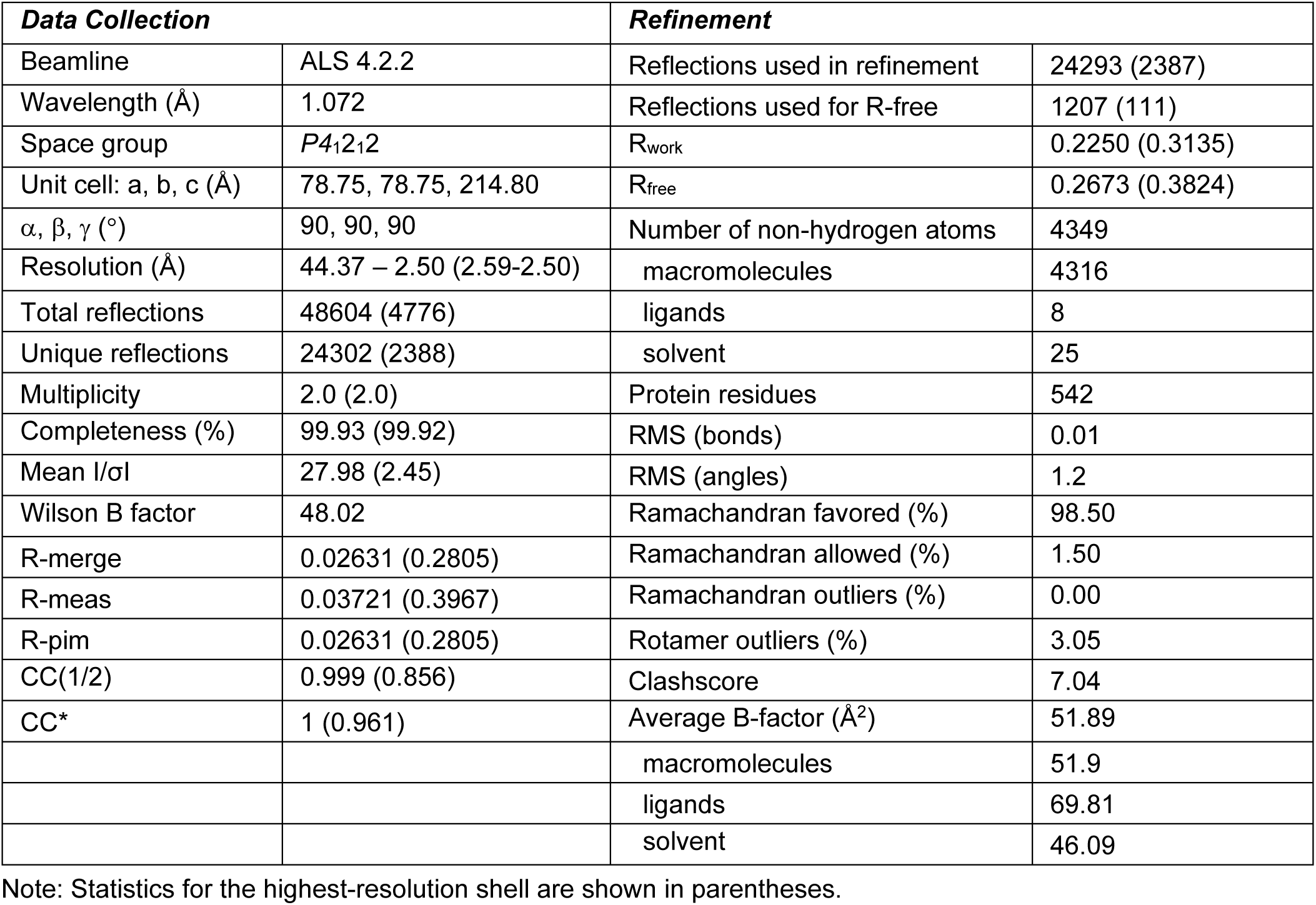
Summary of X-ray data collection and refinement statistics for 7RE5.

| **Data Collection** |  | **Refinement** |  |
| --- | --- | --- | --- |
| Beamline | ALS 4.2.2 | Reflections used in refinement | 24293 (2387) |
| Wavelength (Å) | 1.072 | Reflections used for R-free | 1207 (111) |
| Space group | *P4<sub>1</sub>2<sub>1</sub>2* | R<sub>work</sub> | 0.2250 (0.3135) |
| Unit cell: a, b, c (Å) | 78.75, 78.75, 214.80 | R<sub>free</sub> | 0.2673 (0.3824) |
| $\alpha, \beta, \gamma$ (°) | 90, 90, 90 | Number of non-hydrogen atoms | 4349 |
| Resolution (Å) | 44.37 – 2.50 (2.59-2.50) | macromolecules | 4316 |
| Total reflections | 48604 (4776) | ligands | 8 |
| Unique reflections | 24302 (2388) | solvent | 25 |
| Multiplicity | 2.0 (2.0) | Protein residues | 542 |
| Completeness (%) | 99.93 (99.92) | RMS (bonds) | 0.01 |
| Mean I/σI | 27.98 (2.45) | RMS (angles) | 1.2 |
| Wilson B factor | 48.02 | Ramachandran favored (%) | 98.50 |
| R-merge | 0.02631 (0.2805) | Ramachandran allowed (%) | 1.50 |
| R-meas | 0.03721 (0.3967) | Ramachandran outliers (%) | 0.00 |
| R-pim | 0.02631 (0.2805) | Rotamer outliers (%) | 3.05 |
| CC(1/2) | 0.999 (0.856) | Clashscore | 7.04 |
| CC* | 1 (0.961) | Average B-factor (Å<sup>2</sup>) | 51.89 |
|  |  | macromolecules | 51.9 |
|  |  | ligands | 69.81 |
|  |  | solvent | 46.09 |

Note: Statistics for the highest-resolution shell are shown in parentheses.

The surface of the interface between adjacent Kv2.1 T1 subunits is characterized by complimentary charges (Figure 2A). Electrostatics contribute especially to the N-terminal portion of the T1 domain with a positively charged bulge fitting into a negatively charge concave area on the adjacent subunit. The residues constituting the negatively charged bulge are highly conserved across the T1 family. But the residues constituting the matching positive patch are less conserved so may contribute to the selectivity of T1 domain assembly (Figure 2B).

**Figure 2.**
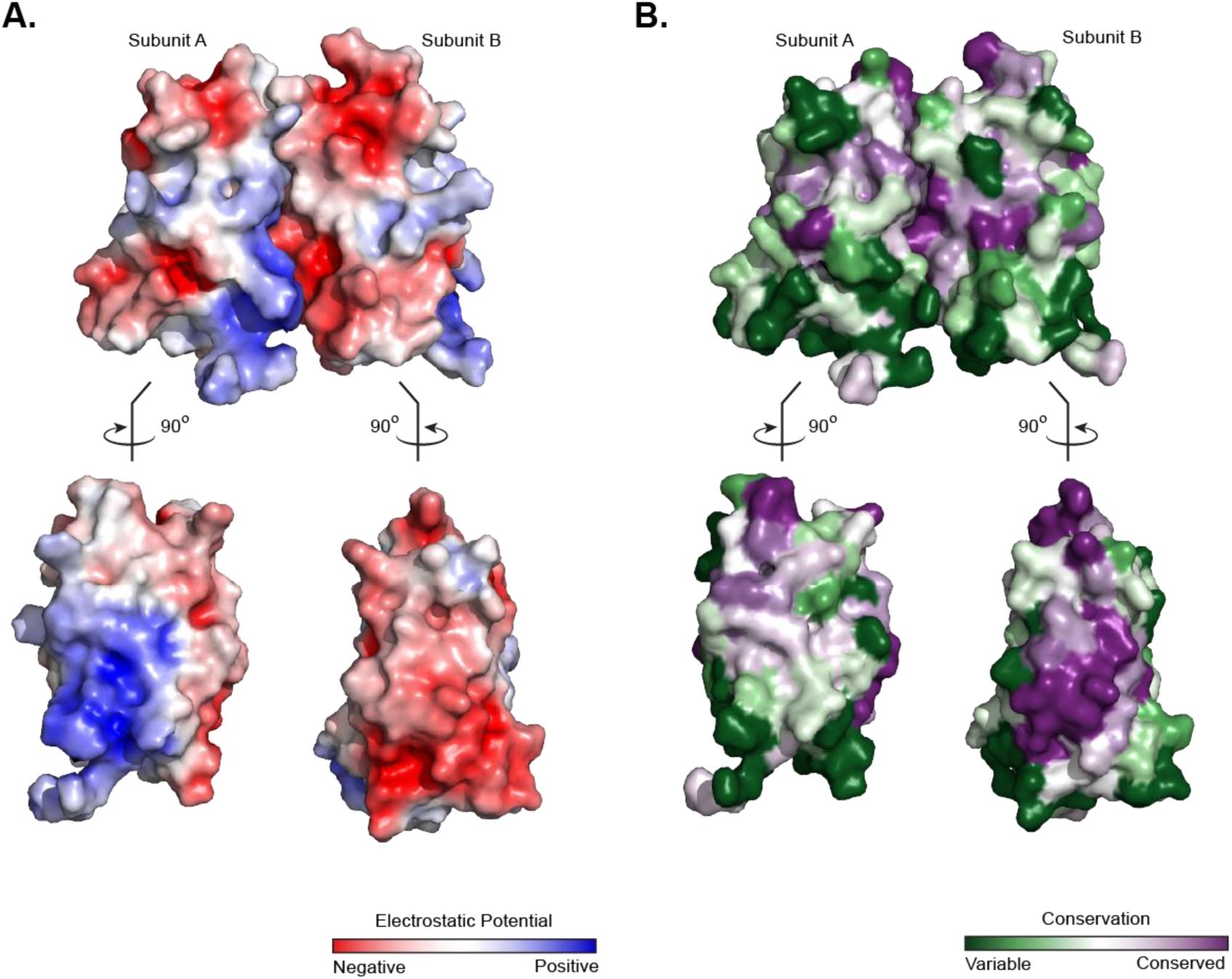
The monomer-monomer interface of Kv2.1 T1 is a conserved charged surface. **(A)** Surface representation of Kv2.1 T1 colored by electrostatics potential calculated using APBS. The gradient is from -5 (red) to +5 (blue) kT/e. *Top*, as observed from side view of X-ray structure in Figure 1B. *Bottom*, subunits are rotated 90° with respect to the X-ray structure. **(B)** Surface representation of Kv2.1 T1 colored by sequence conservation (purple for most conserved and green for least conserved). Conservations scores were calculated using ConSurf. Top and bottom sections are subunits rotated as described for (A).

Two motifs that have been previously implicated in selective higher order assembly of T1 domains are Zn^2+^ binding and a CDD motif (Figure 3A)^12,14-17^. The HX_5_CX_20_CC Zn^2+^ binding motif is found in all Kv2, Kv3, and Kv4 proteins as well as in four of the ten KvS proteins (Kv8.2, Kv6.1, Kv6.2, and Kv6.3) that assemble with Kv2 proteins. This motif, occupied with Zn^2+^, is on the C-terminal end of the T1 domain, which in the context of the full-length protein would be the end closest to the transmembrane domains. In Kv2.1, it is formed by His105, Cys132, and Cys133, with the third cysteine, Cys111, provided by the adjacent subunit (Figure 3B, inset 1). The CDD motif is found in a loop between α3 and β3 on the other end of the subunit interface from zinc (Figure 3B, inset 2). The Asp74 and Asp75 of this motif are in the interface and form a salt bridge with Arg31 and Arg32. The arginines are only conserved in Kv2.1 and Kv2.2. Together these two motifs are well positioned to provide stability to higher-ordered assemblies of Kv2.1 T1 domains.

**Figure 3.**
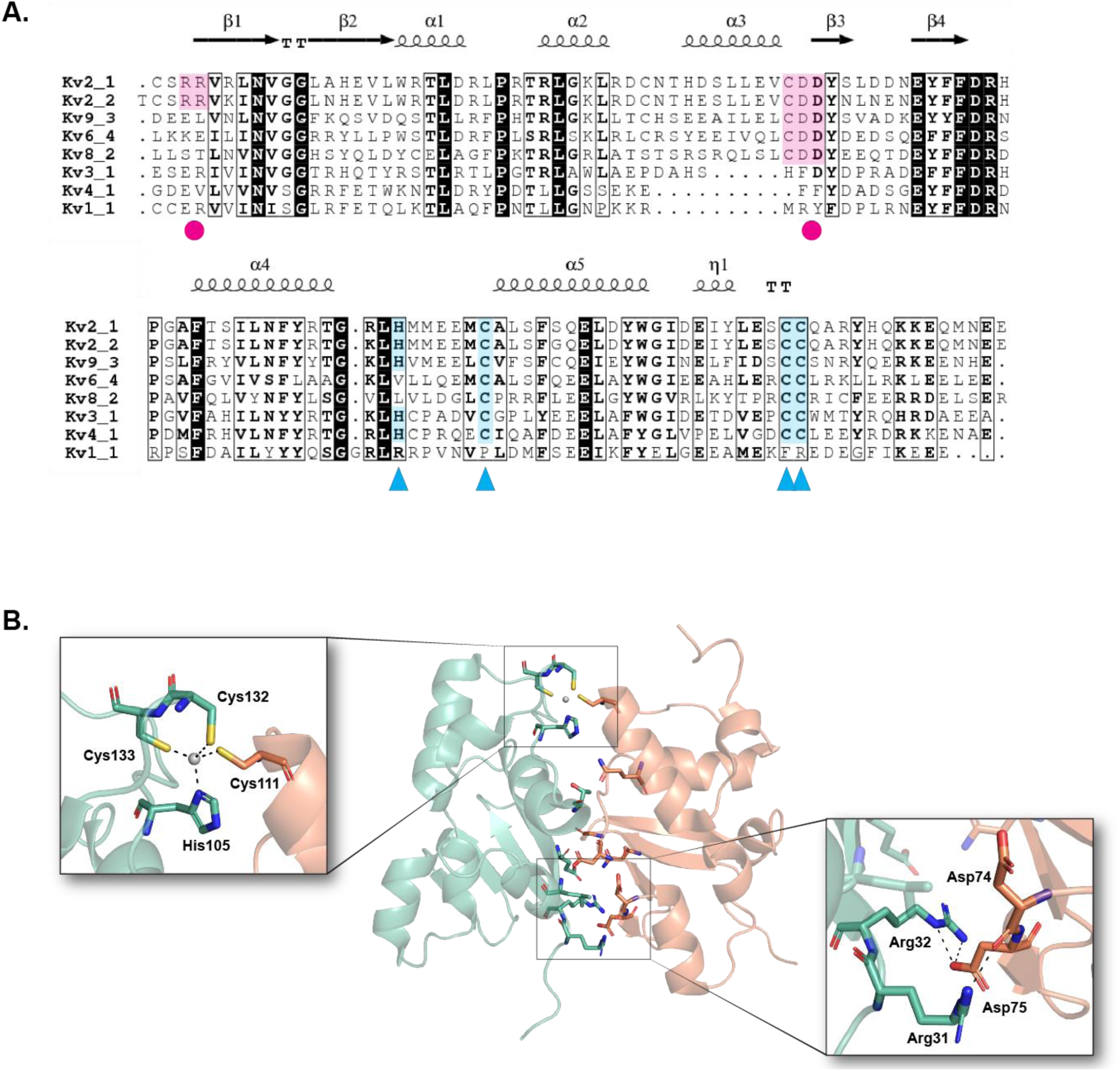
Pentamer formation involves inter-subunit zinc binding and a salt bridge. **(A)** Structure-based multiple sequence alignment of Kv2.1 with KvS (Kv9.3, Kv8.2 and Kv6.4), Kv3.1, Kv4.1, and Kv1.1 T1 domains. Blue triangles below residues indicate the Zn^2+^ coordination motif HX5CX20CC, and magenta circles indicate the negatively charged CDD motif and its interacting positive residues at the N-terminus. Shading of residues in blue or magenta respectively highlight conservation. **(B)** Cartoon representation of Kv2.1 T1 with residues at the subunit interface highlighted as sticks, colors are chain A (green) and chain B (orange). The insets show the interacting residues in the Zn^2+^ coordination sites (*left*) or CDD motif (*right*). Hydrogen bonds are shown in black dashed lines, zinc ion in gray, oxygen in red, sulfide in yellow, nitrogen in dark blue.

### Kv2.1 T1 forms a pentameric conformation in solution

We used multi-angle light scattering and small angle X-ray scattering coupled with size-exclusion chromatography (SEC-MALS-SAXS) as an independent method to determine the stoichiometry of Kv2.1 T1 complexes in solution. The molecular weight of the Kv2.1 T1 monomer is 14.6 kDa, so the theoretical mass of a tetramer is 58.4 kDa while a pentamer is 73 kDa. The absolute molecular mass of Kv2.1 T1 measured by MALS was 71 ± 1 kDa (Figure 4A). The MALS data is most consistent with a pentameric assembly of Kv2.1 T1. From the SAXS data it was determined that the Kv2.1 T1 protein was mono-dispersed (linear in dimensionless Guinier plot) and globular (Gaussian peak in Krakty plot) (Supplemental Figure 2). The experimental scattering curve of Kv2.1 T1 was fit to a tetrameric homology model of Kv2.1 T1 generated using YASARA (based on Kv3.1 T1, PBD: 3KVT); but the fit was very poor, with a χ^2^ = 27. Fitting the experimental scattering curve to the pentameric crystal structure did result in a good fit, χ^2^ =1.7 (Figure 4B). To aid in visualization the calculated surface envelope for Kv2.1 T1 was superimposed with cartoons for the tetrameric homology model or pentameric crystal structure (Figure 4C). The pentameric structure best fit the SAXS data. We also used negative stain electron microscopy (EM) to investigate the oligomeric state of Kv2.1 T1. Top views of ring-shaped particles for Kv2.1 T1 could be seen in the raw electron micrographs, and after 2D class averaging the rings could be more clearly seen to be pentamers (Figure 5). In conclusion, the analysis of Kv2.1 T1 in solution by SEC-MALS-SAXS complimented with direct visualization by EM is consistent with the pentameric assembly revealed by the crystal structure.

**Figure 4.**
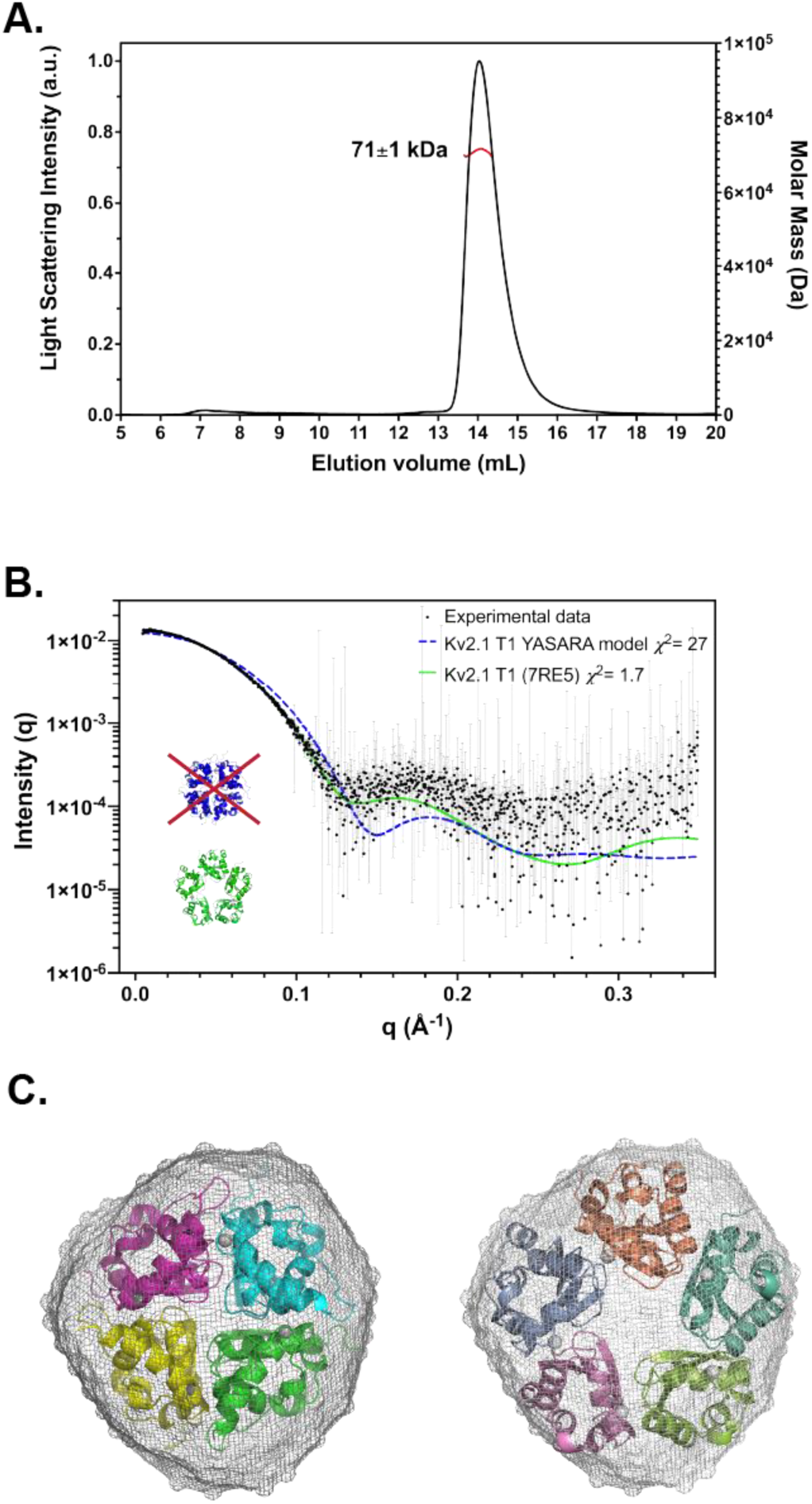
MALS-SAXS analysis of Kv2.1 T1. **(A)** MALS analysis of Kv2.1 T1. The calculated MW value (mean ± SD) taken from the portion of the light scattering peak indicated by the red line. **(B)** Experimental scattering curve of Kv2.1 T1 (black) is compared with the theoretical scattering curves generated by CRYSOL for a tetrameric homology model of Kv2.1 T1 generated using YASARA based on Kv3.1 T1 (PDB: 3KVT), (blue, χ^2^ = 27) or the pentameric Kv2.1 T1 crystal structure (green, χ^2^ =1.7) **(C)** The *ab initio* model of Kv2.1 T1 (shown in surface) was reconstructed using GASBOR and shown is the averaged filtered shape from DAMFILT. The model is superimposed with cartoons for the tetrameric Kv2.1 T1 YASARA homology model (*left*) or the pentameric Kv2.1 T1 crystal structure (PDB: 7RE5) (*right*).

**Figure 5.**
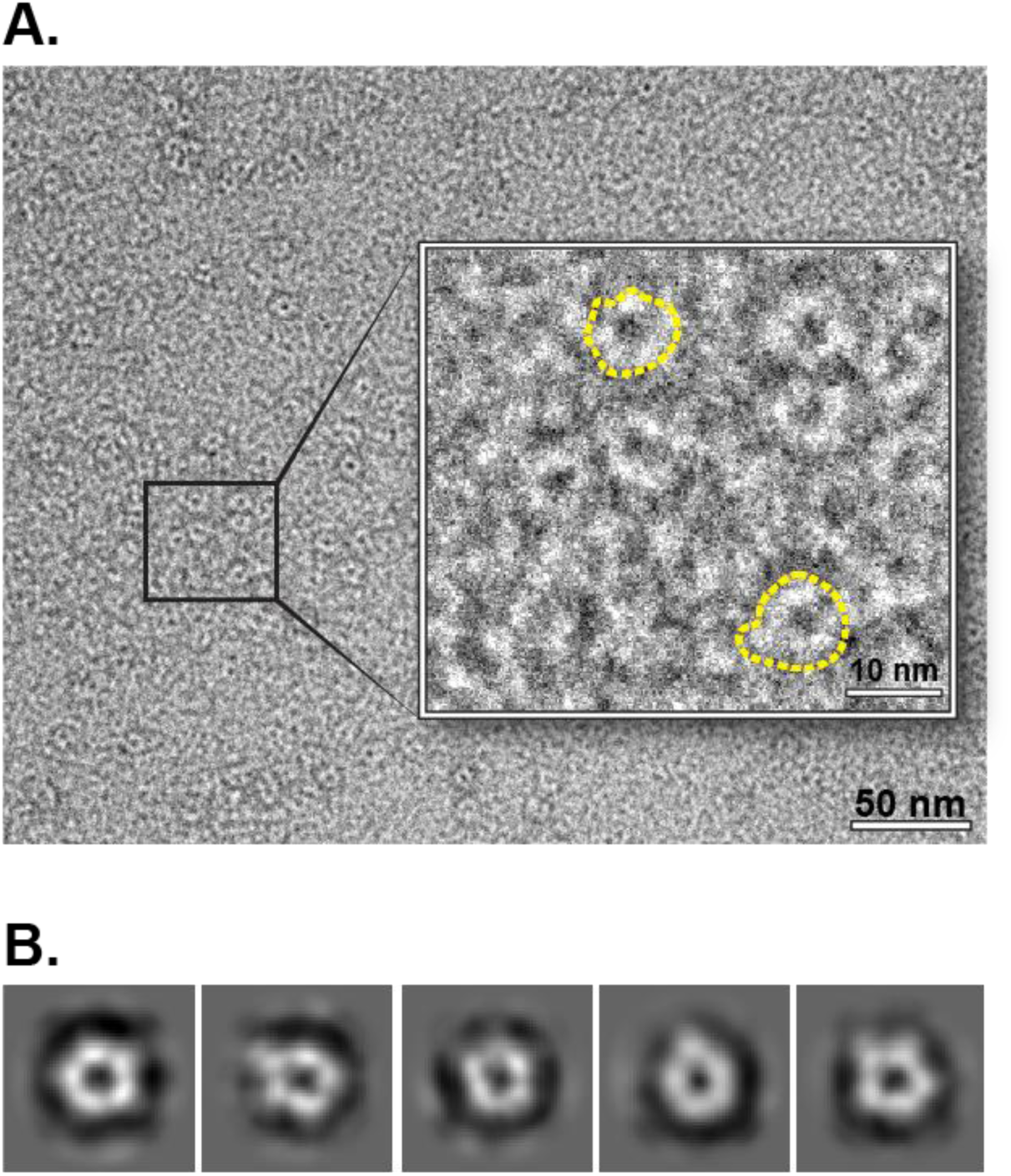
Negative stain transmission electron microscopy of Kv2.1 T1. **(A)** an electron micrograph of Kv2.1 T1 sample negatively stained with 2% (w/v) uranyl acetate and imaged at 100k magnification. *Inset*: increased magnification to aid visualization of individual oligomers, two examples outlined in yellow dashed lines. **(B)** 2D reference-free class averages of particles in top views revealed a pentameric form for Kv2.1 T1.

### The effect of bound Zn^2+^ on the stability of Kv2.1 T1

The role of Zn^2+^ in Kv3 and Kv4 T1 domains has been determined to be critical for the transition from monomers to tetramers as well as protein stability^15,16^. To investigate the effect of bound Zn^2+^ on the assembly and stability of Kv2.1 T1, we used chelation by excess EDTA (1 mM) and assessed the effect that had on the protein using circular dichroism (CD) and dynamic light scattering (DLS) thermal unfolding experiments. The CD spectra calculated for Kv1.1, Kv3.1, and Kv4.1 T1 domains determined using server PDB2CD^20^ was very similar to the experimental spectra obtained for Kv2.1 T1. A strong α-helical signal for Kv2.1 T1 persisted at 10°C after addition of EDTA, implying that Kv2.1 T1 remains in a native fold in this temperature range independent of bound Zn^2+^ (Figure 6A).

**Figure 6.**
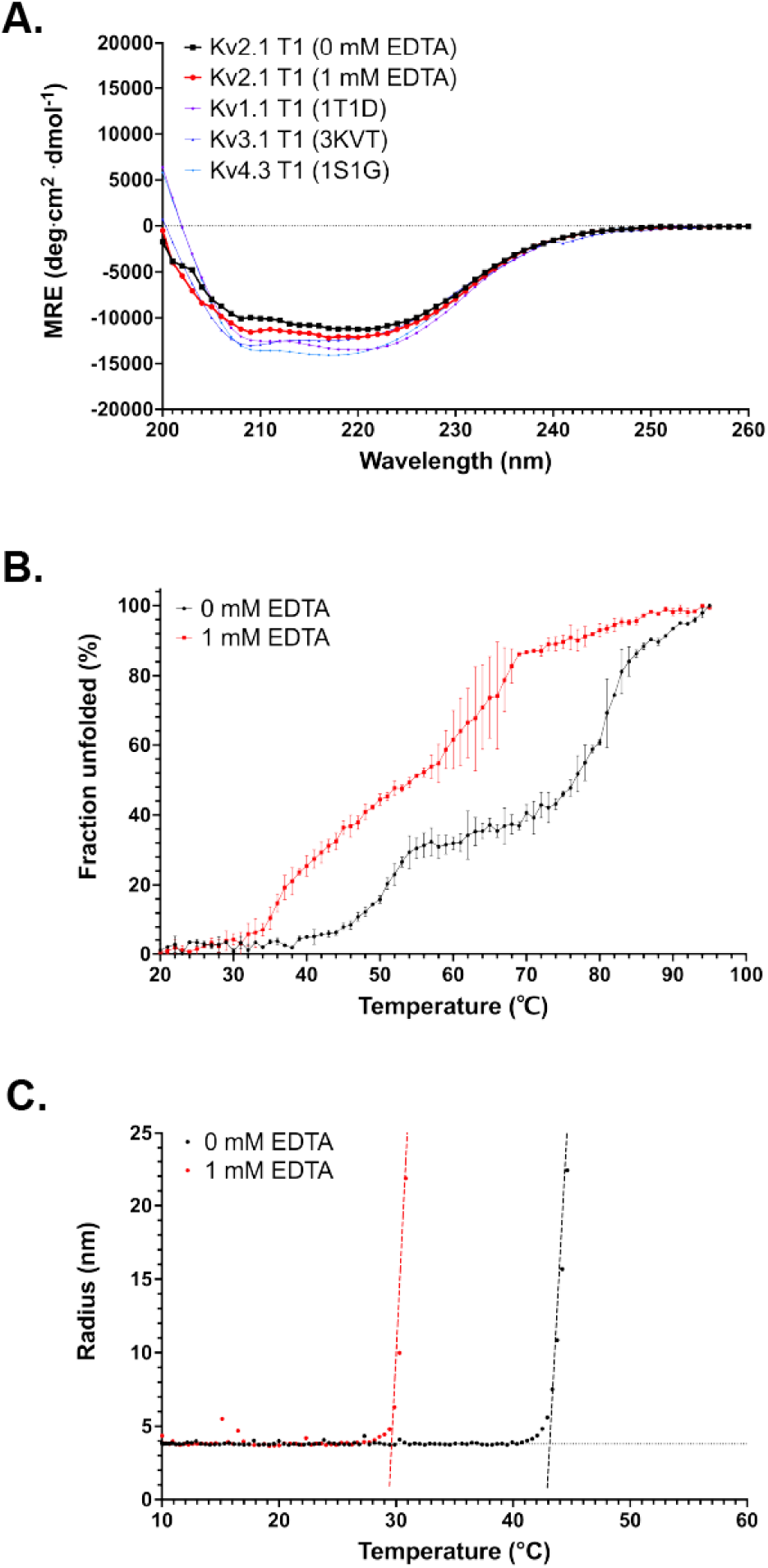
Protein thermal unfolding in the presence of EDTA. **(A)** Comparison of CD spectra of Kv2.1 T1 at 10 °C in the absence ((black) or presence (red) of 1 mM EDTA with the calculated spectra for Kv1.1 (PDB: 1T1D, purple), Kv3.1 (PDB: 3KVT, blue) and Kv4.3 (PDB: 1S1G, light blue) T1 domains. **(B)** Stability analysis by CD of Kv2.1 in solution. The fraction of unfolding extracted from far-UV CD spectra at 222 nm as a function of temperature was plotted. **(C)** Hydrodynamic radius of Kv2.1 T1 monitored by dynamic light scattering are presented as a function of temperature. In B and C, the black data points are from Kv2.1 T1 without EDTA and red data are from Kv2.1 T1 treated with 1 mM EDTA.

The thermal unfolding experiments were monitored by CD at 222 nm at 1 °C intervals from 20-95 °C (Figure 6B). The fraction of unfolded protein (*f*_U_), as a function of temperature exhibited two transitions. The α-helical signal began to diminish at 40 °C and unfolding increased until ∼60 °C; and at ∼80 °C the amount of unfolded protein again increased. In the presence of EDTA the curve was shifted to the left and the differences between the two phases was less striking. To calculate the melting temperature (T_m_, defined as *f*_U_ = 0.5), the fraction of unfolded protein (*f*_U_), as a function of temperature from 10 °C to 70 °C (to encompass the first transition) was fit by a sigmoidal curve. The T_m_ of Kv2.1 T1 was 50 ± 1 °C and in the presence of EDTA, reduced to 38 ± 1 °C. Our results imply that Kv2.1 T1 undergoes a large conformational change at lower temperature in the absence of bound Zn^2+^. Using DLS as a complimentary approach, we monitored the hydrodynamic radius as a function of temperature (10-80°C). The onset transition temperature (or the threshold for when protein unfolding begins; T_m onset_) of Kv2.1 T1, defined by a linear transition plot, was 42.5 ± 0.5 °C. Addition of EDTA reduced the T_m onset_ to 29.5 ± 0.5 °C (Figure 6C). This data supports the CD experiments, and we conclude that Zn^2+^ binding increases the stability of the protein.

## Discussion

Here we report the crystal structure for the T1 domain from Kv2.1. This T1 domain is very similar to the previously described T1 domains from the other Kv subfamilies, except that it was a pentamer. All the previously reported structures for isolated Kv T1 domains are tetramers which is consistent with the tetrameric assembly of full-length Kv subunits into functional channels. Our X-ray structural data is supported by solution analysis of the Kv2.1 T1 domain using SEC-MALS-SAXS and EM imaging.

In the larger family of BTB folds (of which the T1 domain is one class) there is a variety of stoichiometries, from monomers to pentamers^10^. Illustrative to Kv2.1 T1, are the T1-like domains from the Cullin-dependent ubiquitin E3-ligases where the stoichiometry is particularly diverse. In one study, the T1-like domains from SHKBP1, KCTD13, KCTD16, and KCTD17 crystalized as a monomer, tetramer, open pentamer, or closed pentamer respectively even though most of these T1-like domains in the context of full-length native protein are pentamers. In solution, the T1-like domains from SHKBP1 and KCTD13 were predominantly mixtures of monomers and dimers but mixing with the binding partner, Cullin3, induced pentamers^11^.

We propose that for full-length native Kv2.1, the T1 domain assembly state is restrained to a tetramer by interaction with either other parts of the channel or by accessory subunits. In support of this idea, interactions between the cytoplasmic N- and C-termini of Kv2 have been observed^17-19^. Intriguingly, a recent structural study discovered that the orientation of the T1 domain in Kv3 channels is different than in the structures for the Kv1.1-2.1 paddle chimera. The C-terminal most alpha-helix of the Kv3.1a T1 domain is rotated towards and contacts the linker between voltage sensor and pore domains, which may stabilize the pore and contribute to the unique kinetic properties of Kv3 channels^21^. It remains to be determined if a similar arrangement occurs in Kv2 channels.

In terms of accessory subunits, the Kvβ subunit is best characterized for interaction with the Kv1 (shaker) channels and several KChIP proteins serve a similar role to Kvβ for the Kv4 channels^22,23^. Both Kvβ and KChIP proteins make contacts with the T1 domain of the cognate Kv protein and in the case of KChIP can even rescue mutants of Kv4 T1 domains that interfere with tetramerization^24,25^. For Kv2.1 the best characterized interaction proteins are the electrically silent (KvS) alpha subunits^5^ and those mediating the plasma membrane – ER contact sites organized by phosphorylated Kv2.1 in a scaffolding role independent of ion conduction^26-34^. The recent acceleration in solving the structures of ion channels, with and without accessory proteins, by cryo-EM makes it reasonable that a structure of full-length Kv2.1 will be determined and provide information about interactions that influence the T1 domain.

The interface between Kv2.1 T1 subunits in the pentameric ring involves two major features – electrostatics and inter-subunit coordination of zinc. At the N-terminal end of the domain there is a large, highly conserved, patch of surface negative charges that fits into a concavity of positive charges on the adjacent subunit. This is also where the CDD motif is found, a motif recognized for its selective conservation in the Kv2 and KvS proteins^17^. Charge reversal mutants of the aspartates in the CDD motif interfered with assembly Kv2.1. Homology modeling of the Kv2.1 T1 positioned the CDD residues away from the T1 interface so the assembly was thought to have been due to disruption of an interaction with the C-terminus^19^. However, in our structure, the aspartates forming the CDD motif participate in a salt bridge with two arginines at the N-terminus of the T1 thus contribute to stabilizing the T1-T1 interface. Of course, caution must be employed because that salt bridge may be disrupted in a tetramer form of the Kv2.1 T1-T1 interface.

Inter-subunit coordination of zinc for Kv3 and Kv4 T1 domains has been shown to be critical for stability of the protein as well as the assembly of tetramers. Kv4.2 T1 is a particularly interesting case because chelation with EDTA to remove zinc was shown to convert tetramers to monomers which could reform tetramers upon replacement of zinc^15^. For Kv2.1 T1 we obtained two structures, one with full zinc occupancy and one with partial zinc occupancy, in both structures the T1 domain was a pentamer. This indicates that zinc binding alone may not be sufficient to dictate the stoichiometry of the Kv2.1 T1 domain. By using CD and DLS we were able to show that removing zinc by EDTA chelation reduced the stability of the Kv2.1 T1 protein, this latter observation is consistent with the role of zinc binding in stabilizing Kv3 and Kv4 T1 domains.

Our observations of the pentameric Kv2.1 T1-T1 interface has implications for understanding why only T1 domains from Kv2.1 or Kv2.2 can interact with T1 domains from the ten member KvS family. Four of the KvS T1 domains (Kv8.2, Kv6.1, Kv6.2, and Kv6.3) appear to have lost the ability to bind zinc since the histidine in the HX_5_CX_20_CC Zn^2+^ coordination motif is not conserved in those proteins. This alteration would not prevent these T1 domains from providing the cysteine needed to coordinate the Zn^2+^ bound in an adjacent Kv2.1 T1 domain but it would result in only partial zinc occupancy of the tetrameric channels. Since partial zinc occupancy did not prevent crystallization of a Kv2.1 homopentamer, it is possible that partial zinc occupancy is not as deleterious to Kv2.1 containing channels as it is predicted to be for Kv3 or Kv4 containing channels. This could explain why at least four KvS T1 domains do not assemble with Kv3 or Kv4 T1 domains. A feature that impacts all ten KvS T1 domains is the lack of a pair of arginines to form the salt bridge with the aspartates in the CDD motif. If this salt bridge is important in the tetramer as well as the pentamer that we see here, it would imply that the stoichiometry of Kv2: KvS proteins would be most stable in a 2:2 arrangement. Both 2:2 and 3:1 stoichiometries for Kv2: KvS proteins have been observed in transfected cell models but which state predominates in native channels is unknown^35-38^.

In conclusion, we provide the first structural analysis of the human Kv2.1 T1 domain. Since structures for Kv1, Kv3, and Kv4 domains have been solved, we started this work with the expectation of providing a representative T1 domain for the missing channel forming Kv subfamily. We were consequently surprised to find the Kv2.1 T1 domain behaving differently from the others in the stoichiometry that we observed. Future work will need to determine how the Kv2.1 T1 domain is arranged in the context of the full-length channel.

## Experimental procedures

### Cloning, overexpression, and purification of Kv2.1 T1

Human Kv2.1, residues 29-147, corresponding to the T1 domain was cloned using NdeI and BamHI sites in the vector pET28a (Novagen) to express a fusion protein with an N-terminal hexahistidine (6xHis) affinity tag followed by a tobacco etch virus (TEV) cleavage site (ENLYFQG). Transformed *Escherichia coli* (Strain BL21 (DE3)) were grown in LB medium supplemented with 50 ug/mL kanamycin at 37°C to an optical density (OD)_600_ value of 0.7, and 1 mM isopropylthio-β-galactoside (IPTG) was added to induce expression for 15 h at 18°C. Cells were harvested by centrifugation and stored at -80°C. Harvested cells were resuspended in lysis buffer (20 mM Tris-HCl pH 8.0, 150 mM NaCl, 5% glycerol, 0.01% Triton X-100, 20 mM imidazole and 5 mM β-mercaptoethanol (BME)) and supplemented with DNase I and protease inhibitor cocktail (Roche). Cells were disrupted by Emulsiflex C3 (Avestin) to release protein. The His-tagged fusion proteins were purified with a Ni-NTA column (Qiagen), and the elution fraction was incubated with TEV protease for 15-18 h at 4°C to cleave the N-terminal His-tag. The cleaved protein was further purified by a HiLoad® 16/600 Superdex® 200 pg gel filtration column (GE Healthcare) in buffer (20 mM Tris-HCl pH 8.0, 150 mM NaCl, 5% glycerol, 0.01% Triton X-100 and 1 mM DTT). The purified protein was concentrated using Amicon Ultra centrifugal filter devices (Millipore, cut off 10 kDa). The protein samples for dynamic light scattering (DLS) and circular dichroism (CD) measurements were dialyzed into phosphate buffer (20 mM Na_2_HPO_4_ pH 7.4, 150 mM NaCl, 5% glycerol and 1 mM DTT) at 4°C overnight before use.

### Crystallization and X-ray diffraction data collection

Kv2.1 T1 was crystallized using the hanging-drop vapor-diffusion method by mixing 8 mg/ml Kv2.1 T1 in a ratio of 1:1 with the reservoir solution. Initial crystals were obtained at 291 K in 0.2 M magnesium chloride, 15% PEG 400 and 0.1 M sodium HEPES, pH 7.5. Crystals were flash frozen in liquid nitrogen with 30% PEG 400 as cryoprotectant (see PDB: 7SPD). Crystals of Kv2.1 T1 prepared by supplementing the size-exclusion buffer with 50 µM ZnSO_4_ were obtained in 0.2 M Magnesium chloride hexahydrate, 0.1 M Tris pH 8.5, and 3.4 M 1,6-Hexanediol and were flash frozen in liquid nitrogen without additional cryoprotection (see PDB: 7RE5). X-ray diffraction data were collected at 100 K at the Beamline 4.2.2 in Advanced Light Source (Berkeley, CA).

### X-ray diffraction data processing and structure refinement

Diffraction data were indexed, integrated, and scaled using XDS^39^. Initial phase estimates for Kv2.1 T1 were obtained by molecular replacement using Phaser in Phenix^40^, with the structure of Kv3.1 T1 (PDB: 3KVT) as a search model. Refinement was performed using phenix.refine^41^, followed iteratively by manual building using Coot^42^. The final refined structure includes Kv2.1 T1 pentamer with residue 29-133 resolved. Structures were visualized in PyMOL (The PyMOL Molecular Graphics System, Version 2.5 Schrödinger, LLC. Buried surface area was determined using the PISA server^43^. Structural biology applications used in this project were compiled and configured by SBGrid^44^.

### Size exclusion chromatography coupled with multi-angle light scattering and small-angle X-ray scattering (SEC-MALS-SAXS)

The SEC-MALS-SAXS data sets were collected using the BioCAT Beamline 18-ID-D, Advanced Photon Source (Argonne, IL). Samples were centrifuged for 5 min at 13,000 rpm to remove any potential aggregates before injections. 250 μL aliquots containing 1.5 mg/mL of Kv2.1 T1 samples were loaded at a flow rate of 0.5 ml/minute into a 24 mL Superdex 200 Increase 10/300 column on an Agilent 1300 chromatography system. Following elution from the column, the samples were analyzed in line by the UV absorbance detector of the Agilent 1300 chromatography system followed by the DAWN Heleos-II light scattering (LS) and OptiLab T-rEX refractive index detectors in series. Accurate protein molecular weight was determined using the ASTRA software (Wyatt Technology). The elution trajectory was redirected into the SAXS sample flow cell. Scattering data were collected every 1 second using a 0.5 second exposure on a Pilatus 3 1M pixel detector (DECTRIS) covering a q range of 0.0045 < q < 0.35 Å^-1^ (q = 4π/λsinθ, where λ is the wavelength and 2θ is the scattering angle). BioXTAS RAW software was used to collect the SAXS data^45^.

### Small-angle X-ray scattering (SAXS) data processing and modeling

Following data reduction and buffer subtraction, the SAXS data were further analyzed using BioXTAS RAW software. The forward scattering intensity I(0) and the radius of gyration (R_g_) were calculated from the Guinier fit. The normalized Kratky plot and the pair distance distribution plot P(r) and the Porod volume were calculated using the program GNOM embedded in BioXTAS RAW software. The low-resolution *ab initio* bead-based models of proteins were constructed from the experimental data using the program GASBOR^46^. The calculation of theoretical scattering curves for the crystal structure was performed by the program CRYSOL^47^, which also determines the discrepancy (χ^2^ value) between the simulated and experimental scattering curves.

### Circular dichroism (CD) spectroscopy

Far-UV CD spectra of Kv2.1 T1 was acquired using a J-815 spectrometer (JASCO) connected to a Peltier temperature controller. All experiments were performed in duplicate measurements. For each measurement, 300 µl of sample consisting of 20 µM Kv2.1 T1 was added in a 1 mm path length quartz cuvette (Hellma Analytics). The CD ellipticities (θ) were measured from 200 to 260 nm with 1 nm increment, scanning speed set as 50 nm/min and data integration time (D.I.T) set at 2 seconds with a standard sensitivity. Buffers were used as baseline measurements. The final ellipticities were recorded as an average of 4 baseline-corrected scans. The ellipticities (θ) were used to calculate mean residue ellipticity in the formula [θ] = θ/cnl where, c stands for the concentration of protein in moles, n for the number of residues and l for the path length of the cuvette. To examine the effect of Zn^2+^ on Kv2.1 T1 stability, 1 mM EDTA was added to Kv2.1 T1 samples and incubated at 4°C for 12 hours, then the far-UV CD spectra were measured the same as the samples in the absence of EDTA. To detect the thermal unfolding, denaturation experiments were carried out using an automated 1°C incremental temperature ramp in the interval 20–95°C, with a 30 s equilibration time at each measurement step. The thermal unfolding profile of samples were characterized using the mean residue ellipticity minima at 222 nm (θ222) to determine Tm by fitting the Boltzmann sigmoid equation using Prism (GraphPad).

### Dynamic light scattering (DLS)

The thermal ramp stability measurements by DLS were measured in a plastic cuvette with 1 mm path length at various temperatures ranging from 10-80°C, with a ramp rate of 1 °C/minute. Before measurements, the sample was incubated at 10°C for 3 mins. The DLS measurements were averaged from 5 acquisitions of 1 second each. To examine the effect of Zn^2+^ on Kv2.1 T1 stability, 1 mM EDTA was added to Kv2.1 T1 and incubated at 4°C for 12 hours, then the size was measured by DLS and compared to the samples without adding any EDTA. The DLS data were analyzed by Wyatt software and fitted by linear intersection method to determine the Tm onset values.

### Transmission Electron Microscopy

For negative staining, 3 μL protein solution at 0.01 mg/mL was added to a glow-discharged carbon-coated copper grid (Electron Microscopy Science). Grids were stained in 2% (w/v) uranyl acetate (twice for 3 seconds each, followed by a third staining for 25 seconds). Grids were blotted with filter paper (Whatman) to absorb residual solution between each step.

Grids were imaged using a Hitachi HT7800 electron microscope equipped with a tungsten filament and operated at 120 kV. Images were collected at a magnification of 100,000x resulting in a pixel size of 1.93A on the specimen. Only top views of particles were manually picked using cryoSPARC to determine the oligomeric state of Kv2.1 T1^48^. The manually picked particles were classified using 2D reference-free classification and new templates were created for automatic picking to select more particles. Two rounds of 2D reference-free classification was calculated to exclude the “bad” particles and used for data analysis.

## Data Availability

Atomic coordinates and structure factors for Kv2.1 T1 (29-147) with full or partial zinc occupancy have been deposited in the RCSB Protein Data Bank (PDB) with accession codes 7RE5 and 7SPD respectively. SAXS data of Kv2.1 T1 have been deposited in the SASBDB with accession code SASDLJ7.

## Acknowledgments

The authors would like to thank Srinivas Chakravarthy and Jesse Hopkins at BioCAT beamline 18-ID-D (APS) for SEC-MALS-SAXS data collection, Jay Nix at Beamline 4.2.2 (ALS) for X-ray diffraction and fluorescence data collection, and Lokesh Gakhar for helpful discussions.

We acknowledge the use of resources at the Carver College of Medicine’s Protein and Crystallography Facility and Central Microscopy Research Facility at the University of Iowa. This research used resources of the Advanced Photon Source; a U.S. Department of Energy (DOE) Office of Science User Facility operated for the DOE Office of Science by Argonne National Laboratory under Contract No. DE-AC02-06CH11357 and supported in part by a grant from the National Institute of General Medical Sciences of the National Institutes of Health (P30 GM138395 to the Illinois Institute of Technology). This research also used resources of Beamline 4.2.2 of the Advanced Light Source, a U.S. DOE Office of Science User Facility under Contract No. DE-AC02-05CH11231, supported in part by the ALS-ENABLE program funded by the NIH, National Institute of General Medical Sciences (P30 GM124169-01).

## Author contributions

Z.X., N.S. and S.B. conceptualized the work; Z.X., S.K., and N.S. designed and performed experiments and analyzed data. Z.X. and S.B. wrote the manuscript; all authors participated in reviewing and editing the manuscript; N.S. and S.B. supervised the project.

## Conflict of interest

The authors declare that they have no conflicts of interest with the contents of this article.

### Abbreviations and nomenclature

BME: β-mercaptoethanol
BTB: Broad-Complex, Tramtrack and Bric a brac
CD: circular dichroism
DLS: dynamic light scattering
IPTG: isopropylthio-β-galactoside
Kv: voltage-gated potassium channel
SAXS: small angle X-ray scattering
SEC-MALS: size exclusion chromatography coupled with multi-angle light scattering
TEV: Tobacco Etch Virus
T1: Tetramerization domain

## Supplementary Figures

**Supplementary Figure 1.**
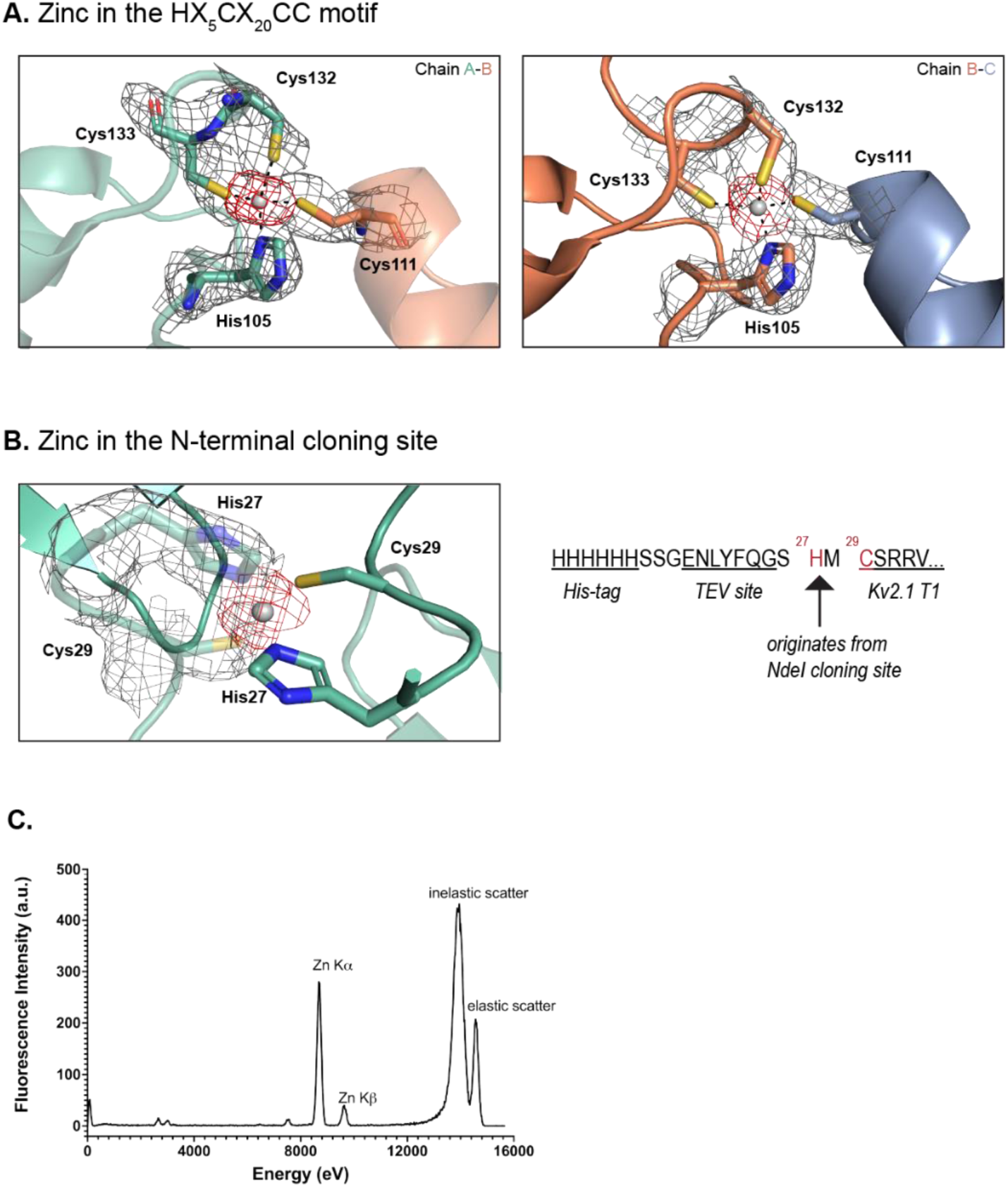
Overview of Zn^2+^ coordination in Kv2.1 T1. **(A)** Tetrahedral coordination of Zn^2+^ by the HX5CX20CC motif. *Left*, the electron densities for Zn^2+^ interaction with His105, Cys132 and Cys133 from chain A and Cys111 from chain B. *Right*, the electron densities for Zn^2+^ interaction with His105, Cys132 and Cys133 from chain B and Cys111 from chain C. Residues are shown in stick representation, zinc ion in gray, sulfide in yellow, nitrogen in dark blue. Coordination bonds are marked with black dashed lines. The gray grid represents the 2mFo – DFc map (1.0σ); red, anomalous map (3.0σ) **(B)** Tetrahedral coordination of Zn^2+^ with His27 (introduced by the NdeI cloning site) and Cys29 from chain A and its symmetry mate in crystal packing. **(C)** Energy spectrum obtained from X-ray fluorescence measurement of the crystal indicates the presence of zinc. The zinc Kα peak is located at 8.6 keV and the zinc Kβ peak is located at 9.6 keV.

**Supplementary Figure 2.**
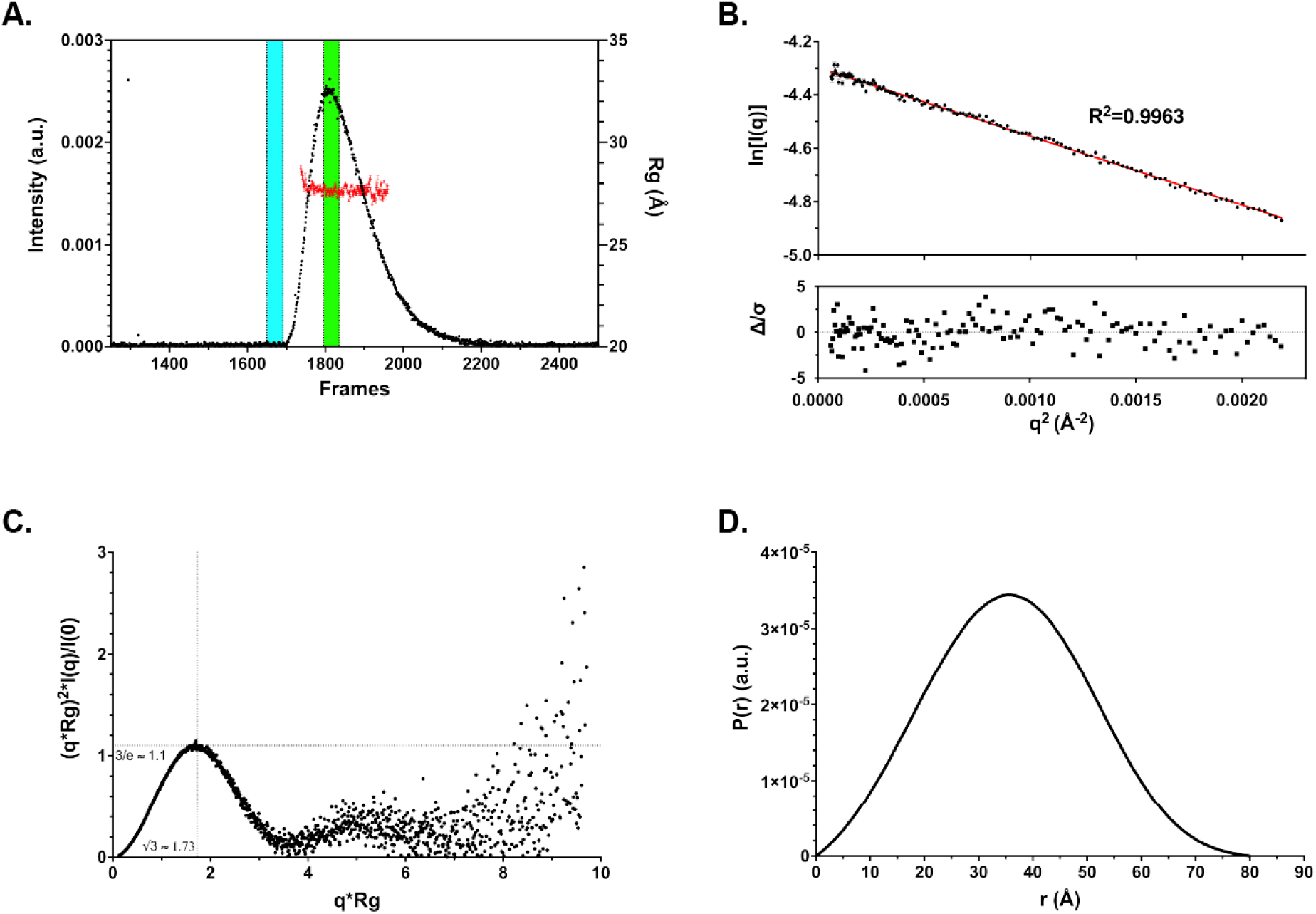
Analysis of Kv2.1 T1 SAXS data quality. **(A)** The total scattering intensity aligns with the Rg plot of Kv2.1 T1 in-line SEC-SAXS frames. The data in the buffer region (cyan) and the sample region (green) were used for buffer subtraction and data analysis. **(B)** The linear low-q region (qRg < 1.3) of the scattering curve was used in Guinier analysis. The Rg of Kv2.1 T1 was 27.7 ± 0.1 Å. **(C)** Rg normalized dimensionless Kratky analysis indicates Kv2.1 T1 is well folded in solution. **(D)** Pair distance distribution P(r) function suggests the Dmax of Kv2.1 T1 was 80 Å.

**Supplemental Table 1.**
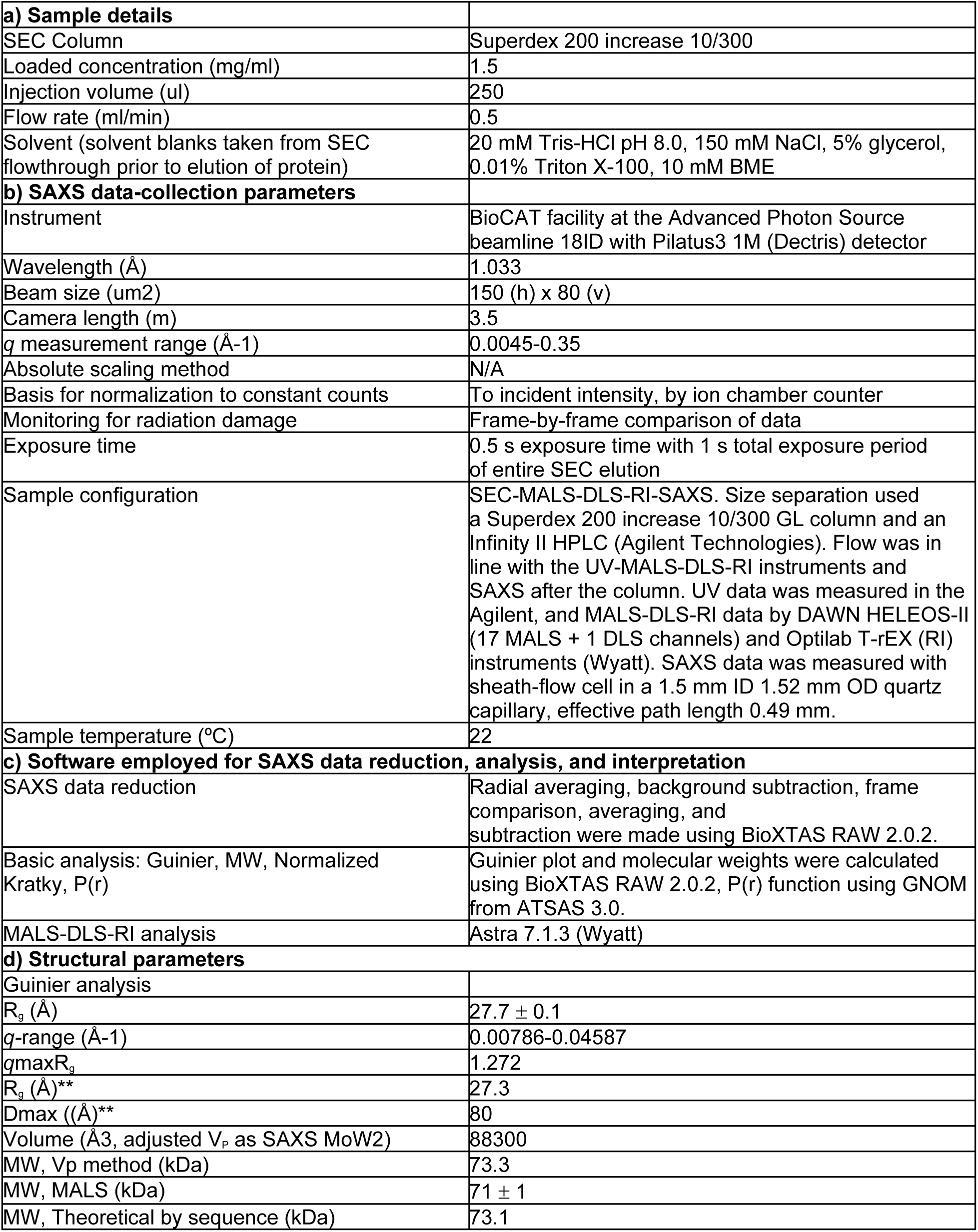
SAXS data collection and analysis parameters

|  |  |
| --- | --- |
| **a) Sample details** |  |
| SEC Column | Superdex 200 increase 10/300 |
| Loaded concentration (mg/ml) | 1.5 |
| Injection volume (ul) | 250 |
| Flow rate (ml/min) | 0.5 |
| Solvent (solvent blanks taken from SEC flowthrough prior to elution of protein) | 20 mM Tris-HCl pH 8.0, 150 mM NaCl, 5% glycerol, 0.01% Triton X-100, 10 mM BME |
| **b) SAXS data-collection parameters** |  |
| Instrument | BioCAT facility at the Advanced Photon Source beamline 18ID with Pilatus3 1M (Dectris) detector |
| Wavelength (Å) | 1.033 |
| Beam size (um<sup>2</sup>) | 150 (h) x 80 (v) |
| Camera length (m) | 3.5 |
| *q* measurement range (Å<sup>-1</sup>) | 0.0045-0.35 |
| Absolute scaling method | N/A |
| Basis for normalization to constant counts | To incident intensity, by ion chamber counter |
| Monitoring for radiation damage | Frame-by-frame comparison of data |
| Exposure time | 0.5 s exposure time with 1 s total exposure period of entire SEC elution |
| Sample configuration | SEC-MALS-DLS-RI-SAXS. Size separation used a Superdex 200 increase 10/300 GL column and an Infinity II HPLC (Agilent Technologies). Flow was in line with the UV-MALS-DLS-RI instruments and SAXS after the column. UV data was measured in the Agilent, and MALS-DLS-RI data by DAWN HELEOS-II (17 MALS + 1 DLS channels) and Optilab T-rEX (RI) instruments (Wyatt). SAXS data was measured with sheath-flow cell in a 1.5 mm ID 1.52 mm OD quartz capillary, effective path length 0.49 mm. |
| Sample temperature (°C) | 22 |
| **c) Software employed for SAXS data reduction, analysis, and interpretation** |  |
| SAXS data reduction | Radial averaging, background subtraction, frame comparison, averaging, and subtraction were made using BioXTAS RAW 2.0.2. |
| Basic analysis: Guinier, MW, Normalized Kratky, P(r) | Guinier plot and molecular weights were calculated using BioXTAS RAW 2.0.2, P(r) function using GNOM from ATSAS 3.0. |
| MALS-DLS-RI analysis | Astra 7.1.3 (Wyatt) |
| **d) Structural parameters** |  |
| Guinier analysis |  |
| *R<sub>g</sub>* (Å) | 27.7 ± 0.1 |
| *q*-range (Å<sup>-1</sup>) | 0.00786-0.04587 |
| *q*<sub>max</sub>*R<sub>g</sub>* | 1.272 |
| *R<sub>g</sub>* (Å)** | 27.3 |
| *D*<sub>max</sub> ((Å)** | 80 |
| Volume (Å<sup>3</sup>, adjusted *V<sub>p</sub>* as SAXS MoW2) | 88300 |
| MW, *V<sub>p</sub>* method (kDa) | 73.3 |
| MW, MALS (kDa) | 71 ± 1 |
| MW, Theoretical by sequence (kDa) | 73.1 |

